# Ethanol-induced conditioned place preference and aversion differentially alter plasticity in the bed nucleus of stria terminalis

**DOI:** 10.1101/519371

**Authors:** Dipanwita Pati, Melanie M. Pina, Thomas L. Kash

## Abstract

Contextual cues associated with drugs of abuse, such as ethanol, can trigger craving and drug seeking behavior. Pavlovian procedures, such as place conditioning, have been widely used to study the rewarding/aversive properties of drugs and the association between environmental cues and drug seeking. Previous research has shown that ethanol as an unconditioned stimulus can induce a strong conditioned place preference (CPP) or aversion (CPA) in rodents. However, the neural mechanisms underlying ethanol induced reward and aversion have not been thoroughly investigated. The bed nucleus of the stria terminalis (BNST), an integral part of the extended amygdala, is engaged by both rewarding and aversive stimuli and plays a role in ethanol seeking behavior. Here, we used ex-vivo slice physiology to probe learning-induced synaptic plasticity in the BNST following ethanol-induced CPP and CPA. Male DBA/2J mice (2-3 months old) were conditioned using previously reported ethanol-induced CPP/CPA procedures. Ethanol-induced CPP was associated with increased neuronal excitability in the ventral BNST (vBNST). Conversely, ethanol-induced CPA resulted in a significant decrease in spontaneous glutamatergic transmission without alterations in GABAergic signaling. Ethanol-CPA also led to significant increase in paired pulse ratio at excitatory synapses, suggestive of a decrease in presynaptic glutamate release. Collectively, these data demonstrate that the vBNST is involved in the modulation of contextual learning associated with both the rewarding and the aversive properties of ethanol in mice.

## Introduction

Alcohol Use Disorder is a highly prevalent chronic disorder characterized by an impaired ability to control alcohol use despite negative health and economic consequences. One of the major obstacles in recovery from addiction is the compulsive drug seeking behavior triggered by environmental contexts and cues contributing to relapse from abstinence (Anton, 1999; Crombag and Shaham, 2002; Krank and Wall, 1990; Monti *et al*, 2002; Zironi *et al*, 2006). This form of Pavlovian learning wherein the rewarding/aversive properties of ethanol become associated with a previously neutral set of environmental stimuli can lead to the maintenance and facilitation of alcohol use disorder. While the focus has been on the rewarding effects of alcohol and how that contributes to alcohol’s addictive potential, a number of studies have also established an equally important role for alcohol’s aversive properties in determining sensitivity to alcohol’s subjective effects (Morean and Corbin, 2010). For example, individuals with family history of alcoholism, are less sensitive to the impairing effects of alcohol (Schuckit, 1994; Schuckit *et al*, 2004). Therefore, alcohol use can be seen as a function of the balance between reward and aversion wherein alcohol’s rewarding properties promote its use while its aversive properties limit its use (Verendeev and Riley, 2012). Thus, understanding the neuroadaptations that underlie these associative memories is integral to the development of better treatment strategies.

A critical neural structure that is widely implicated in associative learning and stress-induced drug seeking is the bed nucleus of the stria terminalis (BNST). BNST is a sexually dimorphic, limbic forebrain region that regulates stress and anxiety states, as well as aversive and reward-related behaviors (for review, see Lebow and Chen, 2016; Vranjkovic *et al*, 2017). It is a complex structure comprising several sub nuclei and heterogenous cell populations (Dong *et al*, 2001a, 2001b, Dong and Swanson, 2006a, 2006b). The anterior commissure divides the BNST into dorsal and ventral (vBNST) divisions. The vast majority of neurons in the BNST are GABAergic in phenotype scattered with a sparse population of glutamatergic cells (Poulin *et al*, 2009; Sun and Cassell, 1993). The BNST receives GABAergic inputs predominantly from the central amygdala (Dong *et al*, 2001b) as well as via intrinsic GABAergic interneurons (Sun and Cassell, 1993). Dense glutamatergic inputs arise from the prefrontal cortex, the basal amygdala, the hippocampus and the insular cortex (Canteras and Swanson, 1992; Cullinan *et al*, 1993; Vertes, 2004). Also, both dorsal and vBNST send primarily GABAergic and glutamatergic projections to the ventral tegmental area (VTA), an area that is known for regulating reward and drug seeking (Dong and Swanson, 2006b, 2006a; Kudo *et al*, 2012). Further, optogenetic stimulation of vBNST-VTA glutamatergic projections resulted in place aversion whereas stimulation of vBNST-VTA GABAergic projections induced place preference (Jennings *et al*, 2013) demonstrating the complex role of BNST in modulating both aversive and appetitive behaviors.

Among the most commonly used behavioral tests in rodents to study cue-induced drug seeking is place conditioning, a classical form of Pavlovian learning that models the rewarding as well as aversive properties of various drugs of abuse (for review see: Bardo and Bevins, 2000; Tzschentke, 1998). Previous literature has shown that distinct tactile cues associated with ethanol can lead to development of conditioned place preference (CPP) in rodents (Ciccocioppo *et al*, 1999; Cunningham and Prather, 1992; Morales *et al*, 2012; Risinger and Oakes, 1996). One of the advantages of CPP is that it can also be used to study contextual association with aversive properties of drugs. Interestingly, the same dose of ethanol can reliably induce either CPP or conditioned place aversion (CPA) dependent on the relative timing of ethanol injection and subsequent exposure to the conditioning stimulus (CS+) (Cunningham *et al*, 1997). Thus, mice injected just before exposure to CS+ developed CPP whereas mice injected right after exposure to the CS+ developed CPA.

Drug-associated contextual cues increase c-Fos immunoreactivity in the BNST (Hill *et al*, 2007; Mahler and Aston-Jones, 2012; Pina *et al*, 2015) supporting a critical role for BNST in driving drug seeking behavior. Disruption of vBNST activity blocked cocaine-induced CPP (Sartor and Aston-Jones, 2012), while chemogenetic inhibition of BNST, and specifically BNST-VTA projections blocked ethanol-induced CPP (Pina *et al*, 2015; Pina and Cunningham, 2017). Several lines of evidence also suggest that neurons in the BNST are susceptible to neuroplastic changes in response to drugs of abuse. For instance, both acute and chronic ethanol exposure altered NMDA receptor function of vBNST neurons (Kash *et al*, 2007, 2009). Nicotine self-administration induced long term potentiation of excitatory synapses in the BNST (Reisiger *et al*, 2014) while self-administration of either cocaine or palatable food enhanced the synaptic strength of excitatory inputs to the vBNST (Dumont *et al*, 2005).

Taken together this emphasizes the importance of BNST in drug-related behavior but very little is known about the neuroadaptations in the BNST following ethanol-induced Pavlovian learning. Hence, our goal was to examine the electrophysiological correlates underlying the neuroplasticity in vBNST following ethanol-induced associative learning using ex vivo patch-clamp electrophysiology. Here, we provide compelling data suggesting ethanol evoked synaptic plasticity in vBNST neurons differ between reward-associated and aversive learning.

## Materials and Methods

### Animals

Male DBA/2J mice aged 2-3 months were obtained from The Jackson Laboratory (Bar Harbor, ME, USA) and housed within UNC animal facilities. Mice were acclimated to the colony for at least one week before being single housed in a temperature and humidity-controlled vivarium with a 12-h light/dark cycle (lights on at 7 am, lights off at 7 pm) with *ad libitum* access to food and water. All experimental procedures were approved by the University of North Carolina Animal and Care and Use Committee’s guidelines.

### Place conditioning paradigm

Mice were conditioned using unbiased, counter-balanced ethanol-induced CPP/CPA procedures as previously reported (Cunningham *et al*, 2006; Pina *et al*, 2015). Briefly, mice were conditioned in a 30 x15 x 15 cm chamber in a sound attenuating box with ~10 lux light intensity between the hours of 9:00 AM-4:00 PM. Chamber floors were divided into two interchangeable halves composed of either grid or hole floor. On day one, mice were allowed to freely explore the chamber for 15 minutes (pretest) to determine initial floor preference. This was followed by four days of conditioning with two 5 min trials per day, where saline (CS−) was administered in the morning and ethanol (CS+; 2 g/kg IP) was administered in the evening. For the CPP procedure, mice were injected with ethanol immediately before 5-min exposure to a distinct tactile cue (grid or hole floor). For the CPA procedure, mice were injected with ethanol immediately following 5-min floor cue exposure. Control mice received saline injections on both CS+ and CS− trials. On the final day, a 30-minute expression test was run to determine preference for the drug-paired floor cue. An additional cohort of mice received either ethanol or saline injections for four days in their home-cages (unpaired) and were included to control for the effects of ethanol and conditioning apparatus exposure.

### Slice electrophysiology

Mice were anesthetized with isoflurane, 1-hour post expression test and were rapidly decapitated. 300 μm thick coronal sections through the BNST were prepared as previously described (Mazzone *et al*, 2016a). Briefly, brains were quickly extracted, and slices were made using a Leica VT 1200s vibratome (Leica Biosystems, IL, USA) in ice-cold, oxygenated sucrose solution containing in mM: 194 sucrose, 20 NaCl, 4.4 KCl, 2 CaCl_2_, 1 MgCl_2_, 1.2 NaH_2_PO_4_, 10 glucose and 26 NaHCO_3_ saturated with 95 % O_2_/5 % CO_2_. Slices were incubated for at least 30 minutes in normal artificial cerebral spinal fluid (ACSF) maintained at 32-35°C that contained in mM: 124 NaCl, 4.4 KCl, 1 NaH_2_PO_4_, 1.2 MgSO_4_, 10 D-glucose, 2 CaCl_2_, and 26 NaHCO_3_, saturated with 95% O_2_/5 % CO_2_. Slices were then transferred to a submerged recording chamber (Warner Instruments, CT, USA) for experimental use. For whole-cell recordings, slices were continuously perfused at a rate of 1.5-2.0 ml/min with oxygenated ACSF maintained at 28±2°C.

Neurons in the ventral BNST were identified using infrared differential interference contrast on a Scientifica Slicescope II (East Sussex, UK). Whole-cell patch clamp recordings were performed using micropipettes pulled from a borosilicate glass capillary tube using a Flaming/Brown electrode puller (Sutter P-97; Sutter Instruments, Novato, California). Electrode tip resistance was between 4 and 6 MΩ. All signals were acquired using an Axon Multiclamp 700B (Molecular Devices, Sunnyvale, CA). Data were sampled at 10 kHz, low-pass filtered at 3 kHz, and analyzed in pClamp 10.6 (Molecular Devices, Sunnyvale, CA). Access resistance was continuously monitored and changes greater than 20% from the initial value were excluded from data analyses. Series resistance was uncompensated and liquid junction potential was not corrected. 2-4 cells were recorded from each animal per set of experiments.

For neuronal excitability experiments, recordings were performed in current clamp mode using a K-gluconate based internal solution (in mM: 135 K-gluconate, 5 NaCl, 2 MgCl_2_, 0.6 EGTA, 4 Na_2_ATP, 0.4 Na_2_GTP, and 10 HEPES, pH adjusted to 7.3 using KOH and volume adjusted to 285–290 mOsm). After allowing cells to stabilize, current was injected to hold the cells at −70 mV and experiments were performed to determine rheobase (the minimum amount of current required to elicit an action potential) and action potential (AP) threshold, followed by spike frequency experiments (number of action potentials fired in response to depolarizing current steps of 10 pA each ranging from 0-80 pA). Parameters related to AP shape, which included AP height, AP duration at half-maximal height (AP half-width), time to fire an AP (AP latency), and afterhyperpolarization (AHP) amplitude were calculated from the first action potential fired during the V-I plot. To assess spontaneous synaptic activity, a cesium methane sulfonate-based intracellular solution (in mM: 135 cesium methanesulfonate, 10 KCl, 1 MgCl_2_, 0.2 EGTA, 4 MgATP, 0.3 Na_2_GTP, 20 phosphocreatine, pH 7.3, 285-290 mOsm with 1mg/ml QX-314) was used. Cells were voltage clamped at −55mV for monitoring spontaneous excitatory post synaptic currents (sEPSCs), and, held at +10mV to isolate spontaneous inhibitory post synaptic currents (sIPSCs). For experiments that required minimal spontaneous synaptic activity in the slice, ACSF was supplemented with tetrodotoxin (TTX; 500 nM), to block voltage-gated sodium channels.

To record evoked currents, a bipolar nichrome electrode was placed in the vBNST dorsal to the recorded neurons. Picrotoxin (25 μM) was added to the ACSF to block both synaptic and extrasynaptic GABA-A receptors to assess post-synaptic glutamate transmission. EPSCs were evoked at 0.167□Hz for 100–150 microseconds using a bipolar Ni-chrome-stimulated electrode controlled by S88X grass stimulators (Astro-Med/Grass technologies) using a cesium gluconate based internal (in mM: 117 D-gluconic acid, 20 HEPES, 0.4 EGTA, 5 TEA, 2 MgCl_2_, 4 Na_2_ATP, 0.4 Na_2_GTP, pH 7.3, 285-290 mOsm with 1mg/ml QX-314). Cells were held at −70 mV to record AMPA receptor mediated currents and held at +40 mV to record NMDA receptor mediated currents. The NMDA current at +40 mV was measured at least 50 ms after the onset of an evoked response to avoid contamination with AMPA current. To assess putative changes in presynaptic release probability, paired pulse ratio (amplitude of EPSC2/ EPSC1) was recorded at 50 ms inter stimulus interval while the cells were voltage clamped at −70 mV. Similar set of experiments were conducted to assess plasticity at GABAergic terminals using a cesium chloride based internal (in mM: 130 CsCl, 10 HEPES, 2 Mg-ATP, 0.2 Na_2_GTP, pH adjusted to 7.3 using CsOH and volume adjusted to 285–290 mOsm) in the presence of kynurenic acid (3mM) to block ionotropic glutamate receptors. Since eIPSCs have slower kinetics, PPR was calculated as delta amplitude of IPSC2 divided by amplitude of IPSC1.

### Drugs

All chemicals used for slice electrophysiology were obtained from either Tocris Bioscience (Minneapolis, USA) or Abcam (Cambridge, UK). For CPP/CPA experiments ethanol (95%) was prepared 20% v/v in 0.9% saline solution and administered intraperitoneally at a dose of 2 g/kg in a 12.5 ml/kg volume.

### Data and statistical analysis

Place conditioning data were analyzed by comparing percent time spent within ethanol-paired chamber and saline-paired chamber following expression test. For differences between two groups, a standard unpaired t-test was used, and Welch’s correction was applied in cases where the groups had unequal variance. Differences in various electrophysiological measures were compared between saline and ethanol-treated groups. Repeated measures ANOVAs (treatment X current injection) were used to assess between group differences in the spike numbers fired across a range of current steps. Sidak’s multiple comparisons post hoc tests were used to analyze significant ANOVA terms. Correlation analyses were computed using Pearson’s correlation coefficient. All data are expressed as mean ± SEM. P-values ≤ 0.05 were considered significant. All statistical analysis was performed using Graph Pad Prism v.7 (La Jolla, CA, USA).

#### Exclusion criteria

Whole cell recordings were not performed in EtOH-CPP mice that showed aversion (<50% time on CS+) during the expression test (n=6), or EtOH-CPA mice that showed preference (>50% time on the CS+) during the expression test (n=2). For various electrophysiological measures, Grubbs’ test was used to detect outliers where appropriate.

## Results

### Expression of ethanol-induced CPP and CPA in adult male DBA/2J mice

To examine synaptic plasticity in the BNST following ethanol-induced associative learning, we trained mice using the classical Pavlovian model of place conditioning. Mice were conditioned to associate one floor, but not another, with ethanol. The CPP model was used to assess the rewarding properties of ethanol. In this model, mice were injected with ethanol (2g/kg) immediately before a 5 min CS exposure (Fig. 1A). Prior to conditioning, a 15 min pretest was included to determine initial bias (Fig. 1B-C). No initial bias for the CS+ side was observed between Sal (n=24 mice) and EtOH (n=24 mice) groups [t(46) =1.178, p = 0.9133]. Also, there was no difference in the mean velocity (Fig. 1C) between the two groups during pretest [t(46) =0.8545, p = 0.3973]. After four days of conditioning, mice were given free access to both the conditioning floors during a drug-free test day to determine the expression of ethanol preference. Consistent with previous reports (Pina *et al*, 2015), mice treated with ethanol showed increased preference for the CS+ floor (Fig. 1D; [t(46) = 3.117, p=0.0031]). This was also accompanied with a significant decrease in mean velocity in the EtOH group which is indicative of enhanced expression of CPP (Cunningham, 1995; Vezina and Stewart, 1987) (Fig. 1E; [t(46) = 2.112, p = 0.0402]).

**Figure 1:**
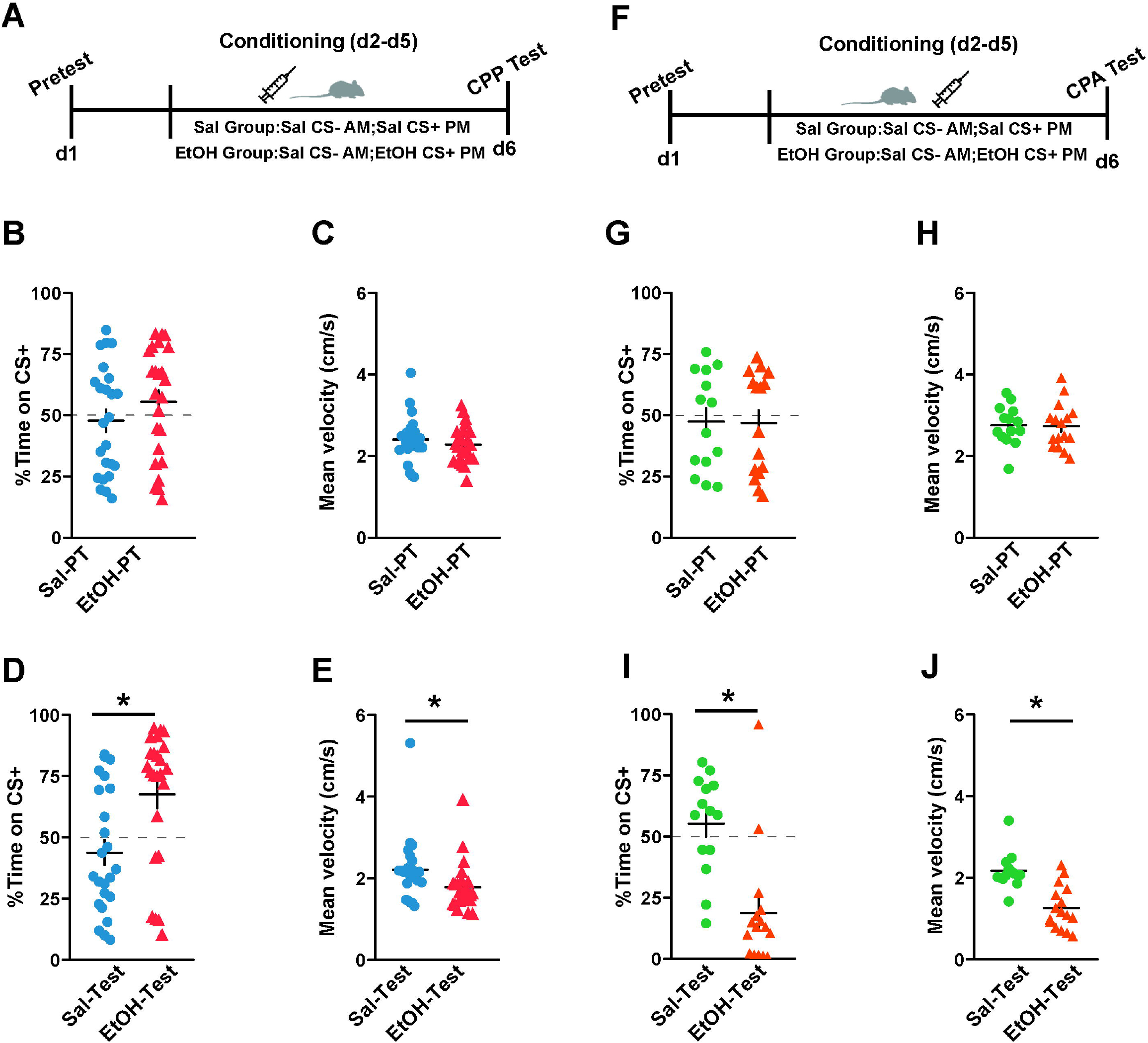
Ethanol-induced Pavlovian learning in male DBA2/J mice. **A)** Experimental timeline for ethanol-induced CPP paradigm. **B-C)** No difference in preference for CS+ side was observed between mice assigned to saline group (n=24) and mice assigned to ethanol group (n=24). There was also no difference in mean velocity between the two groups. **D-E)** Following four days of conditioning with two 5 min trials per day, mice injected with ethanol showed significant preference for CS+ side on a drug-free test day when compared to saline treated mice. Post conditioning, mice in ethanol-paired group showed a significant reduction in mean velocity compared to saline-paired mice. **F)** Experimental timeline for ethanol-induced CPA paradigm. **G-H)** Average pretest data showing mice assigned to saline group (n=14) didn’t differ in their preference for CS+ side when compared to ethanol group (n=16). No difference in mean velocity between the two groups were observed. **I-J)** Mice injected with ethanol immediately after 5 min exposure to a distinct cue (grid or hole floor) expressed significant aversion for CS+ side on Test day and also showed a significant reduction in mean velocity compared to saline-paired mice following ethanol-induced aversion. Data expressed as Mean ± SEM.*p<0.05.

In a separate cohort of mice, we injected the same dose of ethanol immediately after a 5 min CS exposure to induce CPA (Fig. 1F). Similar to the CPP paradigm, we found no difference in the time spent on grid and hole floors during the pretest between the Sal (n=14 mice) and EtOH (n=16) groups (Fig. 1G; [t(28) = 0.0752, p = 0.9406)]), and there were no differences in basal locomotion between treatment groups (Fig. 1H; [t(28) = 0.1513, p = 0.8808]). On the test day, mice treated with ethanol showed a significant aversion to the CS+ floor (Fig. 1I; [t(28) = 4.449, p=0.0001]). This conditioned aversion was associated with a reduction in mean velocity in the EtOH mice (Fig. 1J; [t(28) = 5.055, p<0.0001]). Together, this shows that we can reliably obtain both ethanol-induced CPP and ethanol-induced CPA in male DBA/2J mice.

### Increase in intrinsic excitability of vBNST neurons following ethanol-induced CPP is mediated through inhibition of inward-rectifying potassium channels

To characterize synaptic plasticity in BNST neurons following ethanol conditioning, mice were sacrificed an hour after the expression test for ex vivo patch clamp electrophysiological recordings. Recordings were performed in the vBNST, which has a dense population of projection neurons interspersed with interneurons and has been implicated in addictive behaviors. Intrinsic properties of vBNST neurons were calculated from data generated in voltage clamp using brief voltage steps from −70 to −80 mV. We found that the average input resistance in the saline-CPP group was 495.1 ± 46.27 MΩ and in the ethanol-CPP group it was 626.4 ± 72.03 MΩ ([t(39.22)=1.535, p=0.1329]; unpaired t-test with Welch’s correction). The average capacitance was generally <30 pF and didn’t differ between the two groups ([t(46)=1.266, p=0.2118]). In order to evaluate the effect of ethanol-induced place preference, the intrinsic excitability of vBNST neurons was assessed through rheobase (the minimum current required to elicit an action potential), action potential threshold and the number of action potentials fired across a range of current steps (0-80 pA, at an increment of 10 pA). All the measurements were recorded at −70 mV in current clamp mode to account for variability in resting membrane potentials across different neurons. EtOH-CPP did not alter the resting membrane potential (RMP) between the two groups (Fig. 2B; n=19 cells from 7 saline mice; n=15 cells from 6 ethanol-treated mice; [t(32) =1.360, p=0.1835]) but significantly reduced both the rheobase in the ethanol-treated mice (Fig. 2C; n=24 cells from 9 mice in CPP group; Sal mice (n=24 cells from 7 mice); [t(36.59) = 2.745, p = 0.0093]; unpaired t-test with Welch’s correction) and the action potential threshold (Fig. 2E; [t(46) = 3.152, p = 0.0029]). The reduction in average rheobase in the EtOH-CPP mice showed a significant negative correlation with the average time spent on the CS+ (Fig. 2D; r^2^=0.5183; p=0.0287; Pearson correlation coefficient). Significant between-group differences were also observed in the number of action potentials generated across a range of current injections, a revealed by repeated measures two-way ANOVA (Fig. 2F-G; [F(7,322) = 5.1899, p<0.0001] for group x current interaction; [F(1,46) = 23.66, p<0.0001] for main effect of group; [F(7,322) = 68.09, p<0.0001] for main effect of current). Together, these results indicate that ethanol-induced CPP results in hyperexcitability of vBNST neurons.

**Figure 2:**
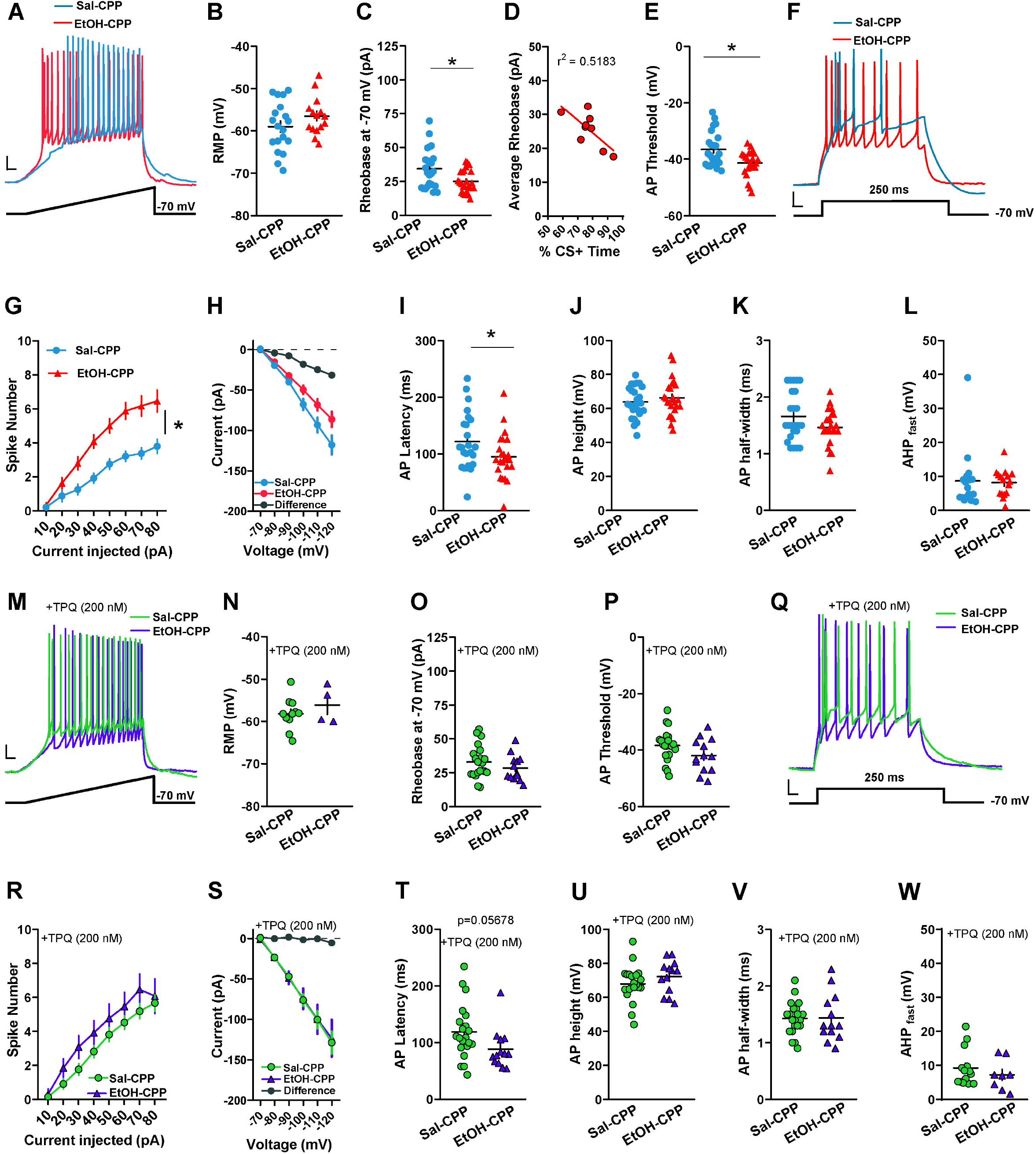
Ethanol-induced CPP increases excitability in vBNST neurons through inhibition of inward rectifying potassium channel. **A)** Representative data obtained from vBNST neurons in Sal and EtOH mice in response to a 120 pA/s current ramp while injecting a constant current to hold the cell at −70 mV. The minimum current required to elicit an action potential (rheobase) was reduced in EtOH mice (n=24 cells from 9 mice) compared to Sal (**C**; n=24 cells from 7 mice) without any changes **(B)** in resting membrane potential (RMP). **D**) There was a negative correlation between average rheobase for each animal and percentage time spent on CS+ side (r^2^=0.5183; p=0.0287). **E)** We also observed a significant decrease in action potential threshold. **F-G)** Representative traces of a neuron in Sal and EtOH group, respectively firing action potentials in response to a step protocol of increased current steps of 10 pA/250 ms. There was a significant increase in number of spikes in response to a graded current injection in the EtOH group. **H)** Current-voltage relationship of vBNST neurons indicates a reduction in an inwardly-rectifying current in the EtOH-treated mice. **I-L)** Ethanol-induced CPP also altered the action potential kinetics resulting in decreased latency **(I)** and a trend toward decreased action potential half-width (**K**; p=0.0802) in EtOH mice. There was no difference between EtOH and Sal mice groups in average action potential height **(J)** and fast after hyperpolarization potential **(L)**. In a different cohort of mice, excitability experiments **(M-W)** were conducted in the presence of Tertiapin-Q (TPQ, 200 nM) to block inward-rectifying potassium channels. No change in RMP **(N)** was observed between the two groups in the presence of TPQ. Representative data **(M)** obtained from vBNST neurons in the showing no change in rheobase between saline-treated (n=21 cells from 10 mice) and ethanol-treated (n= 13 cells from 5 mice) groups **(O)**. **P)** There was still a trend (p=0.0893) toward decrease in action potential threshold in the EtOH group compared to the Sal group. **Q-R)** There was no significant difference in number of spikes in response to a graded current injection between the two groups in the presence of TPQ. **S)** There was no difference in current-voltage relationship of vBNST neurons between the two groups. **T-W)** Ethanol-induced CPP decreased AP latency **(T)** but not average action potential height **(U)**, action potential half-width **(V)** or fast after hyperpolarization potential **(W)** in the presence of TPQ. Scale bar (10 mV, 250 ms), lines in dot plots represent mean ± SEM. *p<0.05.

Accumulating evidence suggests that various ion channels (such as K+ channels) can alter the kinetics of action potential and modulate intrinsic excitability. Plasticity of intrinsic excitability is critical for synaptic integration and learning processes and by modulating various ion channels drugs of abuse may contribute to alterations in neuronal firing (Kourrich *et al*, 2015). Therefore, we dissected the action potential shape in different components (Fig. 2I-L), for example, spike latency, fast after-hyperpolarization (AHP fast), action potential height and half-width. Interestingly, we observed a decrease in AP latency in the EtOH group (Fig. 2I; [t(46) = 2.027, p= 0.0485]) which was accompanied by a trend toward decrease in spike half-width (Fig. 2J; [t(46) = 1.789, p = 0.0802]) without affecting either spike height (Fig. 2K; [t(46) =0.8195, p=0.4167] or fast AHP (Fig. 2L; n=17 cells in Sal group and n=16 in EtOH group; [t(22.66)=0.2292, p=0.8208]; unpaired t-test with Welch’s correction). To further investigate the contribution of ion channels on excitability, we performed voltage clamp experiments to evaluate the relationship between holding current and command potential between −70 mV and −120 mV with 10 mV current steps (Fig. 2H). A two-way repeated measures ANOVA revealed a main effect of group [F1,47=4.206, p=0.0459], along with a significant group × voltage interaction [F(5,235)=4.608, p=0.0005] and main effect of voltage [F(5,235)=178.8, p<0.0001]). Consistent with this observation, the difference between these two I-V plots, highlights that neuronal excitability following ethanol-induced reward learning is potentially associated with decreased activation of an inwardly rectifying current that reverses near the equilibrium potential for potassium.

To directly test the hypothesis that hyperexcitability of vBNST neurons in the ethanol-treated mice involves inactivation of inward-rectifying potassium channels, we ran a separate cohort of mice through the CPP paradigm and measured various excitability related parameters in slices extracted from both ethanol-treated and saline-treated mice in the presence of Tertiapin-Q (TPQ, 200 nM) to selectively block inward rectifier potassium channels (Fig. M-W). There was no difference between average input resistance across both the groups (441.2 ± 59.72 MΩ in saline group and 475 ± 48.37 MΩ in ethanol-CPP mice [t(32)=0.3970, p=0.6940]). In the presence of TPQ, we did not observe any differences in either rheobase (Fig. 2O; n= 21 cells from 10 saline-treated mice and n=13 cells from 5 ethanol-treated mice; [t(32)=1.135, p=0.2647]) or in the number of action potentials fired across a range of current steps (Fig. 2R; [F(7, 224) = 0.7305, p = 0.6463] for group x current interaction; [F(1, 32) = 2.046, p = 0.1623] for main effect of group; [F(7, 224) = 77.54, p<0.0001] for main effect of current; RM measures two-way ANOVA) between the two groups. Further, in the presence of TPQ, no difference between the two I-V plots (Fig. 2S; [F(5,160) = 0.0760, p = 0.9958] for group x voltage interaction; [F(1, 32) = 0.0041, p = 0.9489] for main effect of group; [F(5,160) = 96.22, p<0.0001] for main effect of voltage; RM measures two-way ANOVA) was observed further suggesting the involvement of inward-rectifying potassium channels. Interestingly, even though blocking of inward rectifier potassium channels prevented the increase in excitability of vBNST neurons in the EtOH mice, there was a trend toward reduction in the action potential threshold (Fig. 2P; [t(32)=1.752, p=0.0893]) compared to Sal mice. Analysis of action potential shape revealed that while there was no difference in spike height (Fig. 2U; [t(32) =1.167, p=0.2518], spike half-width (Fig. 2V; [t(32) = 0.0807, p=0.9361]) or fast AHP (Fig. 2W; n=14 cells in Sal group and n=8 in EtOH group; [t(20)=0.8809, p=0.3889]), the reduction in AP latency in the EtOH-treated mice still persisted in the presence of TPQ (Fig. 2T; [F(32)=1.968, p=0.0578]). This suggests that alterations in AP threshold and latency observed in EtOH-treated mice are not mediated by inward-rectifying potassium channels but through some other ion channel mechanism. Collectively, these data provide a potential mechanism by which ethanol-induced place preference produces a functional increase in the excitability of vBNST neurons.

### No change in intrinsic excitability of vBNST neurons following ethanol-induced CPA

We next asked whether ethanol-induced conditioned aversion might also modulate excitability of vBNST neurons. Mice that showed conditioned aversion to ethanol (n=7) were similar to saline treated mice (n=8) with regards to RMP (Fig. 3A; n=8 cells from Sal mice and n=11 cells from EtOH mice; [t(17) =0.6776, p=0.5072]), rheobase (Fig. 3B; n=16 cells from Sal group and n=25 cells from EtOH treated mice; [t(20.97) =1.165, p=0.2572]), action potential threshold (Fig. 3C; [t(39)=0.5266, p=0.6014]) and in the number of action potentials fired across a range of current steps (Fig. 2D;[F(7, 273) = 0.1993, p = 0.9854] for group x current interaction; [F(1,39) = 0.1809, p=0.6729] for main effect of group; [F(7,273) = 86.16, p<0.0001] for main effect of current; RM measures two-way ANOVA). We included a third cohort of mice that received either ethanol (2g/kg) or saline injections in their home cage for four consecutive days to account for the pharmacological effects of ethanol injection in the absence of any explicit conditioning. Ex vivo patch clamp recordings were conducted from vBNST neurons on the fifth day. Similar to EtOH-CPA mice, no group differences were evident between unpaired Sal (n=4) or EtOH mice (n=4) across various parameters: RMP (Fig. 3E; n=11 cells from Sal and n=8 cells from EtOH mice; [t(17) =0.846, p=0.41], rheobase (Fig. 3F; n=14 cells in Sal group and n=14 in EtOH group; [t(26) =0.9912, p=0.3307], AP threshold (Fig. 3G; [t(26)=0.1424; p=0.8879]) and in V-I plot (Fig. 3H; [F(7,168) =0.2370, p=0.97] for group x current interaction; [F(1,24) =0.1946, p=0.6630] for main effect of group; [F(7,168) = 41.46, p< 0.0001] for main effect of current; RM measures two-way ANOVA). Collectively, these results suggest that ethanol-induced increases in neuronal firing is selective for associative reward learning.

**Figure 3:**
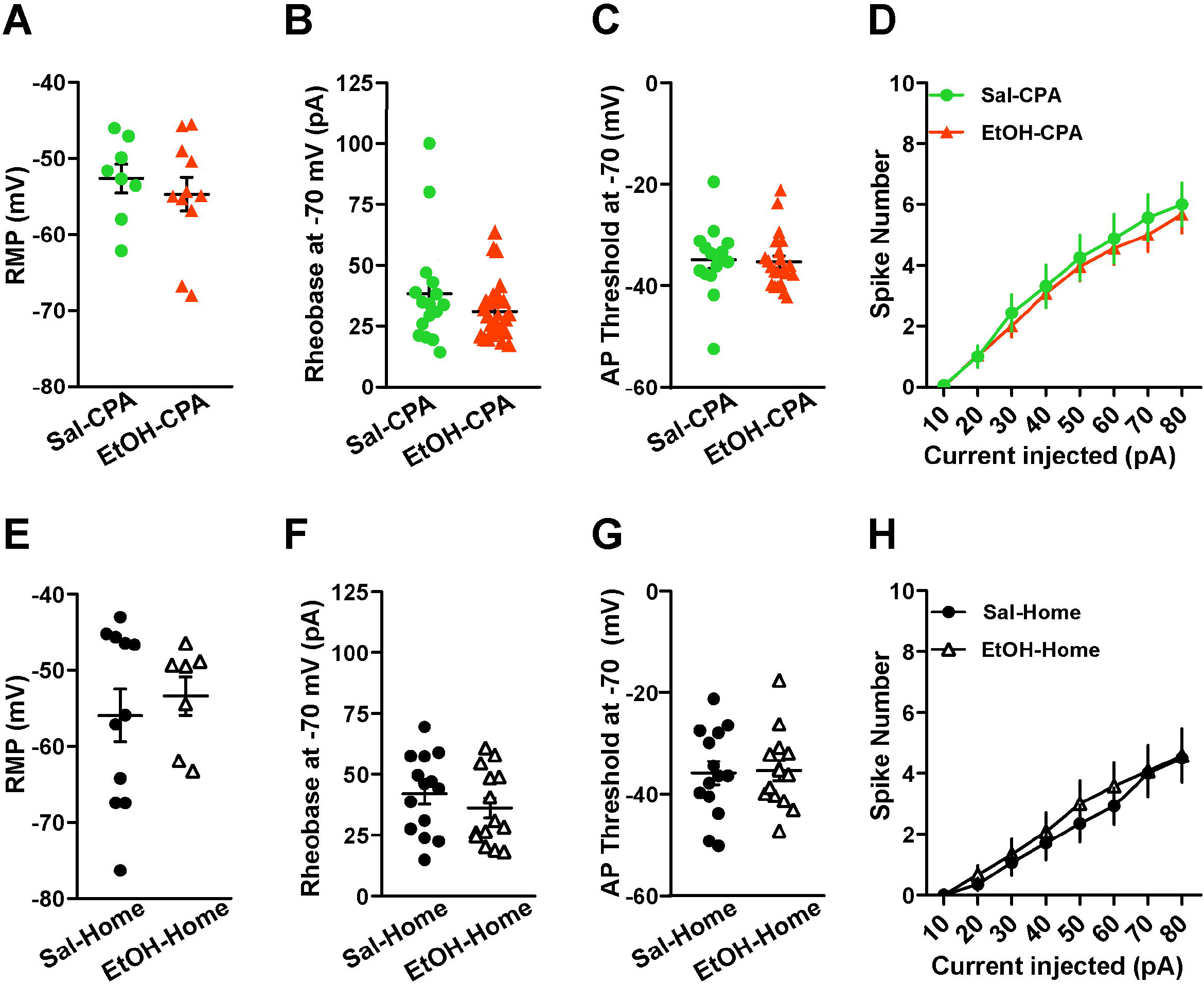
No changes in excitability in vBNST neurons following either ethanol-induced CPA or unpaired-ethanol injections. EtOH mice (n=7 mice) did not have significant difference in RMP **(A)**, rheobase **(B)**, action potential threshold **(C)** or in the number of spikes in response to current steps **(D)** compared to Sal mice (n=8 mice) following ethanol-induced CPA. Similarly, mice that received ethanol injections in their home cages (n=4 mice) didn’t vary significantly across various excitability measures **(E-H)** from mice that received home cage saline injections (n=4 mice). Lines in dot plots represent mean ± SEM.

### Decrease in spontaneous glutamatergic neurotransmission to the vBNST neurons associated with ethanol-induced CPA, but not ethanol-induced CPP

Ethanol-induced Pavlovian conditioning can influence excitatory drive on vBNST neurons by altering spontaneous excitatory or inhibitory synaptic transmission. To evaluate this, we used a Cs-methanesulfonate based internal solution (see methods) to measure both spontaneous excitatory postsynaptic currents (sEPSCs) when voltage clamped at −55 mV, and spontaneous inhibitory postsynaptic currents (sIPSCs) in the same cells when voltage clamped at +10 mV (Fig. 4). The advantage of using this approach is the ability to approximate excitatory-inhibitory (E-I) drive on individual neurons. Ethanol-induced conditioned preference had no significant effect on either sEPSC frequency (Fig. 4B; n=19 cells from 6 Sal-paired mice and n=24 cells from 6 EtOH-paired mice; [t(41)=0.2763, p=0.7837]) or amplitude (Fig. 4C; [t(27.33) =0.6128, p=0.5451]; unpaired t-test with Welch’s correction). Similarly, no change was observed in the frequency of sIPSC (Fig. 4B; [t(37)=1.691, p=0.0993]; unpaired t-test with Welch’s correction) or amplitude of sIPSC (Fig. 4C; [t(41)=0.3218, p= 0.7493]).

**Figure 4:**
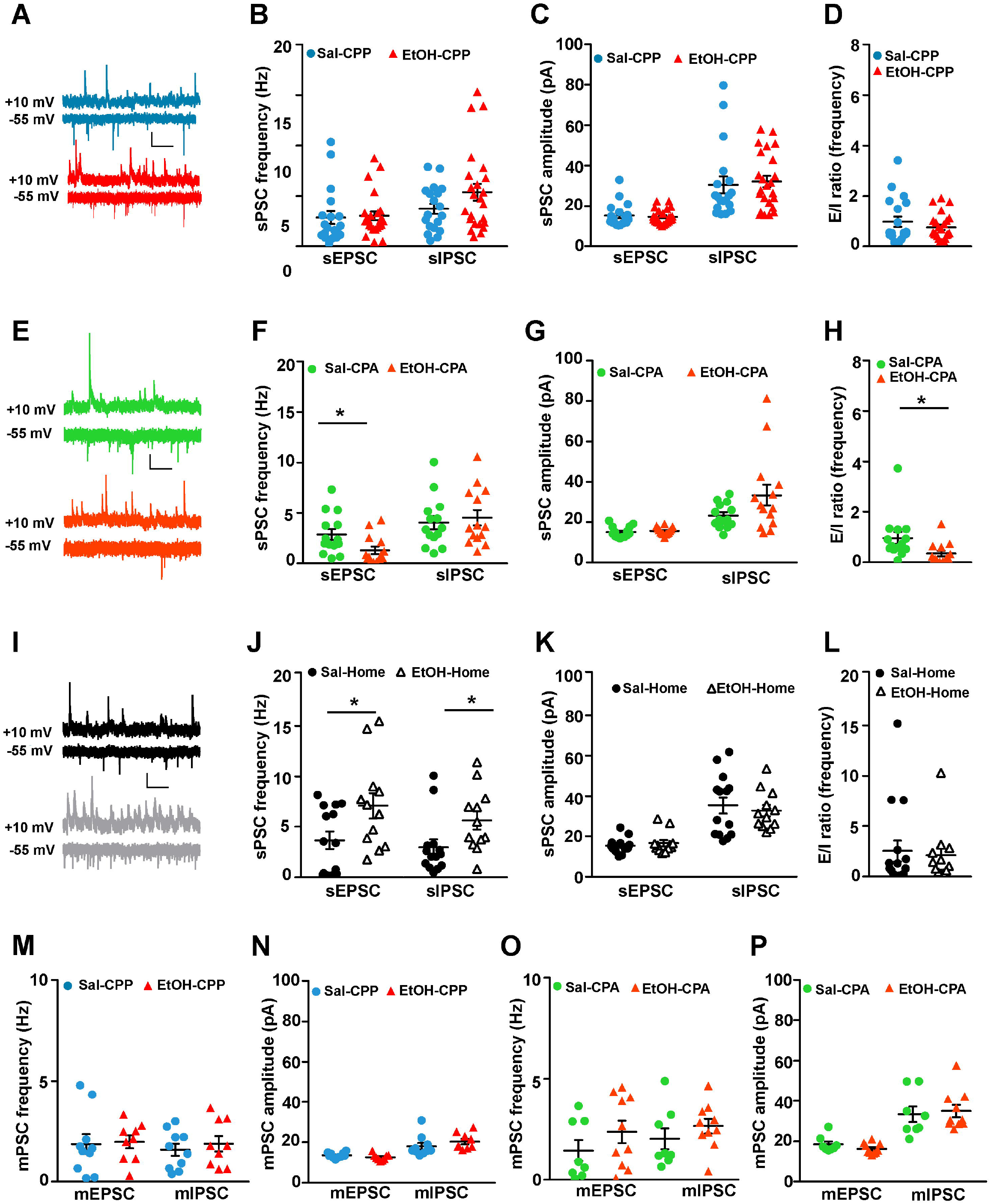
Ethanol-induced CPA decreases spontaneous excitatory synaptic transmission in vBNST neurons. **A)** Representative traces of spontaneous postsynaptic currents (sPSCs) from Sal-CPP (n=6 mice; 19 cells) and EtOH-CPP mice (n=6; 24 cells). There were no differences in sEPSC or sIPSC parameters (frequency, amplitude and excitation-inhibition ratio) of vBNST neurons **(B-D)** between the two groups. **E)** Representative data from Sal-CPA (n=5) and EtOH-CPA mice (n=5) showing a reduced excitatory drive on vBNST neurons following ethanol-induced aversion. There was a significant decrease in sEPSC frequency in the EtOH mice (**F**; n=14 cells) without any alterations in sIPSC frequency when compared to Sal mice (n=14 cells). No significant differences were observed in sEPSC or sIPSC amplitude **(G)**. The excitation-inhibition ratio of vBNST neurons in EtOH mice was also significantly lower than Sal mice **(H)**. **I)** Representative traces of sPSCs from Sal-Home (n=6; 15 cells) and EtOH-Home mice (n=6; 12 cells) indicating an overall increase in both GABAergic and glutamatergic transmission following exposure to ethanol. There was a significant increase in both sEPSC and sIPSC frequency **(J)** without any alterations in amplitude **(K)** when compared to Sal mice. The excitation-inhibition ratio of vBNST neurons didn’t vary significantly between Sal mice and EtOH mice **(L)**. **M-P)** There were no differences in miniature postsynaptic currents (mPSCs) following either ethanol-induced CPP (n= 4 mice in each group) or CPA (n=2 in sal group; n=4 EtOH group). mPSCs were isolated by adding TTX (500 nM) to block action-potential mediated synaptic transmission. Scale bar (20 pA, 500 ms). Lines in dot plots represent mean ± SEM. *p<0.05.

Interestingly, we found that ethanol-induced conditioned aversion decreased spontaneous excitatory synaptic drive onto vBNST neurons. Specifically, in slices extracted from mice that received ethanol we observed a significant reduction in sEPSC frequency (Fig. 4E-H; n=of 14 cells from 5 mice in each group; [t(26)=2.419, p=0.0229]) without any changes in sIPSC frequency [t(26)=0.4901, p=0.6282], resulting in overall decrease in E-I drive on vBNST neurons [t(18.33)=2.375, p=0.0287; unpaired t-test with Welch’s correction]. Mean amplitude were unaltered in both the groups [t(26)=0.6409, p=0.5272] for sEPSCs while there was a trend toward increase in sIPSC amplitudes [t(15.5)=1.893, p=0.0771] that failed to reach significance.

Spontaneous post synaptic currents (sPSCs) recorded from mice that received ethanol injections in their home cages revealed an overall increase in spontaneous synaptic transmission when compared to control mice (Fig. 4I-L; n=6 mice in each group). There was a robust increase in sEPSC frequency (n=14 cells in Sal and n=12 cells in EtOH group, respectively; [t(24)=2.275, p=0.0321]) without any change in amplitude. This was also accompanied with an increase in frequency of inhibitory transmission [t(24)=2.249, p=0.0340] but not amplitude. Though, we didn’t see any net changes in the synaptic drive onto vBNST neurons in the two groups [t(24)=0.4394, p=0.6643].

To investigate the role of activity-independent transmission in ethanol-induced place conditioning, we bath applied TTX (500 nM) to isolate action potential-independent miniature neurotransmission. There were no effects of either EtOH-CPP (n=10 cells from 4 Sal mice and n=9 cells from 4 EtOH mice) or EtOH-CPA (n=8 cells from 2 Sal mice and n=10 cells from 4 EtOH mice) in any of the mPSC measures (Fig. 4M-P). Collectively, these data reveal a strong reduction in spontaneous excitatory drive on vBNST neurons selectively in mice that demonstrated ethanol-induced place aversion.

### Alterations in short-term presynaptic glutamate plasticity following ethanol-induced CPA, but not ethanol-induced CPP

Previous results suggest that there may be a presynaptic inhibition of excitatory transmission on vBNST neurons in EtOH-CPA mice that is network activity-dependent. To further address this, in the next set of experiments, we performed electrically evoked synaptic recordings (Fig. 5) from vBNST neurons. First, we evaluated synaptic facilitation, a form of short-term plasticity (Zucker and Regehr, 2002). We bath applied picrotoxin (25 μM) to block GABA-A receptors and isolate glutamatergic EPSCs. Then, we evoked pairs of excitatory post-synaptic currents by delivering electrical stimulation 50 ms apart. This allows us to measure paired pulse ratio (PPR; amplitude of pulse 2/amplitude of pulse 1). Alterations in PPR are reflective of potential changes the probability of presynaptic neurotransmitter release. We found that only ethanol-induced place aversion caused significant increase in PPR, suggesting a putative decrease in neurotransmitter release probability from glutamatergic inputs to vBNST when compared to control animals (Fig. 5B; n=17 cells from 6 Sal mice and n=22 cells from 7 EtOH mice; [t(37)=2.163, p=0.0371]). EtOH-induced CPP did not alter glutamatergic PPR relative to saline controls (Fig. 5A; n=21 cells from 6 Sal mice and n=15 cells from 5 EtOH treated mice; [t(34)=0.5527, p=0.5841]). Furthermore, unpaired ethanol injection also did not affect PPR ratio at glutamatergic terminals (Fig. 5C; n=15 cells from 4 mice in each group; [t(28)=0.3086, p=0.7599]).

**Figure 5:**
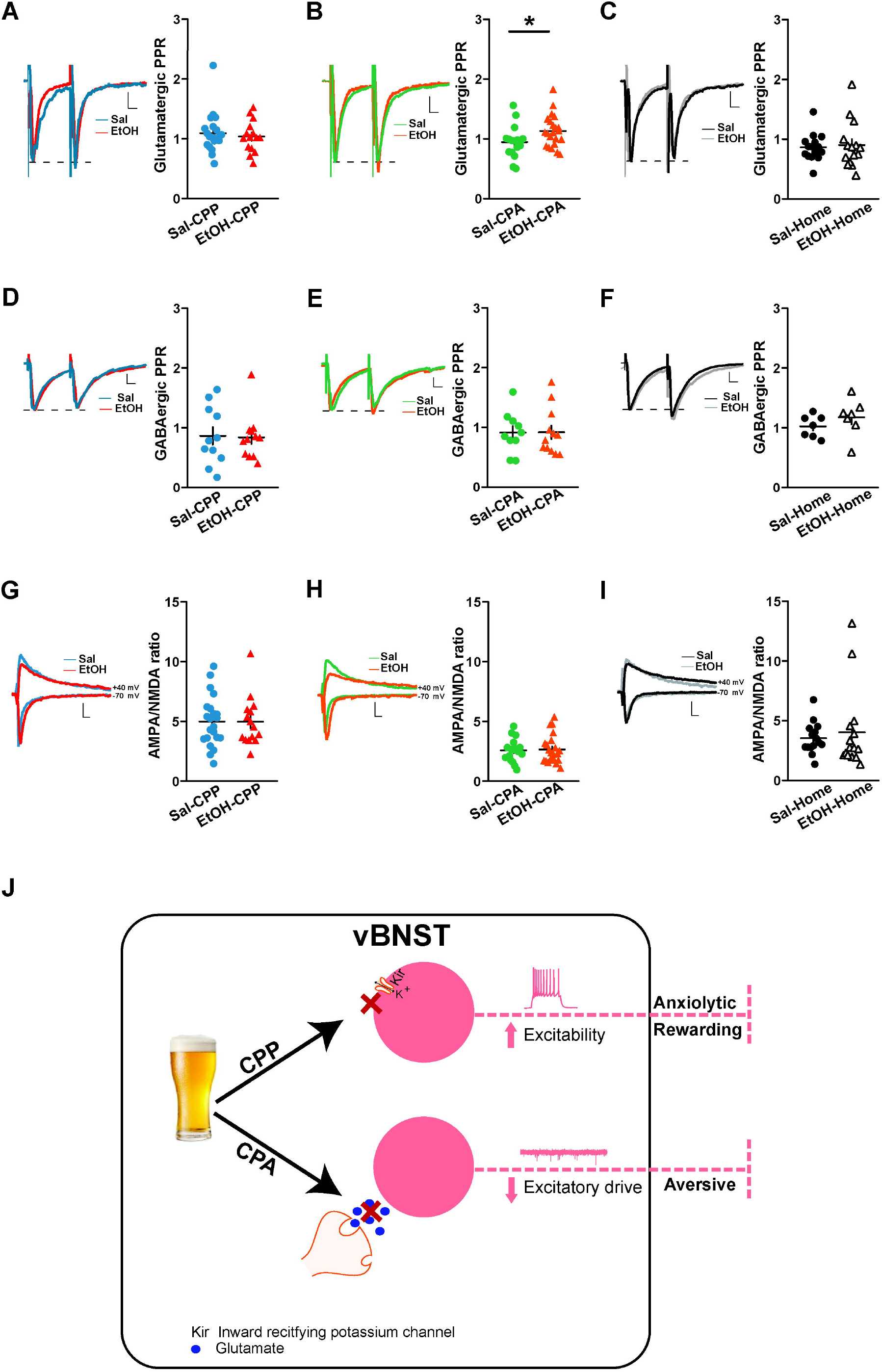
Ethanol-induced CPA selectively decreases presynaptic release probability at glutamatergic terminals without altering AMPA/NMDA ratio. **A)** Representative traces of paired pulse ratios (PPR with 50 ms inter stimulus interval; amplitude of pulse 2/ amplitude of pulse 1) at glutamatergic synapses following ethanol-induced CPP. All traces are normalized to first pulse for each graph. Scale bar (50ms). No effect of ethanol-induced CPP on glutamatergic PPR was observed in the two groups (Sal: n=21 cells from 6 mice; EtOH: n=15 cells from 5 mice). **B)** Representative traces of glutamatergic PPR following ethanol-induced CPA. There was a significant increase in PPR in the EtOH mice (n=17 cells from 7 mice) when compared to Sal mice (n=22 cells from 6 mice), indicative of potential decrease in presynaptic release probability. **C)** No changes were observed in glutamatergic PPR following home cage exposure to ethanol (n=15 cells from 4 mice in each group). **D-F)** Representative traces of paired pulse ratios (PPR with 50 ms inter stimulus interval; amplitude of pulse 2/ amplitude of pulse 1) at GABAergic synapses. Traces are normalized to first pulse for each graph. Scale bar (100 ms). No changes were observed in PPR at GABAergic terminals following exposure to ethanol when compared to their respective saline controls. (n=11 cells from 3 mice in Sal group and n=12 cells from 3 mice for EtOH-CPP experiments; n=10 cells from 3 mice in Sal-CPA and n=12 cells from 4 mice in EtOH-CPA group; n=7 cells from 2 mice in each group for unpaired ethanol exposure). G-I) Representative traces of AMPA/NMDA ratio (AMPA current measured at −70 mV; NMDA current measured at +40 mV after a delay of 50 ms) following exposure to ethanol-induced CPP. All traces are normalized to NMDA for each graph. Scale bar (100 ms). No effect of ethanol-induced CPP on AMPA/NMDA ratio was observed in the two groups (**G**; n=15 cells from 4 Sal mice; n=14 cells from 5 EtOH mice). **H)** Ethanol-induced CPA didn’t affect the AMPA/NMDA ratios between the Sal mice and EtOH mice (n=17 cells from 6 Sal mice; n=22 cells from 7 EtOH mice). **I)** Similarly, home cage exposure to ethanol (n=15 cells from 4 mice) didn’t significantly alter the AMPA/NMDA ratios when compared to the saline control mice (n=15 cells from 4 mice). **J)** Model depiction of ethanol-induced neuroadaptations in the vBNST following Pavlovian learning. Ethanol-CPP increases neuronal excitability by blocking inward-rectifying potassium channels on putative VTA-projection neurons which can be rewarding while ethanol-CPA reduces excitatory drive on vBNST neurons via modulation of presynaptic release probability at glutamatergic synapses which can potentially drive aversion. Lines in dot plots represent mean ± SEM. *p<0.05.

We also evaluated potential changes in presynaptic release probability at GABAergic terminals by evoking GABAergic IPSCs in presence of kynurenic acid (3 mM) to block ionotropic glutamatergic receptors (Fig. 5D-F). Ethanol exposure in both the conditioning paradigms ([t(21)=0.1399, p=0.8901] in CPP; [t(20)=0.0432, p=0.9649] in CPA) and under home cage conditions ([t(12)=1.107, p=0.2900]) did not affect the inhibitory PPR.

Thus, ethanol-induced place aversion selectively modulated short-term plasticity by putatively altering presynaptic glutamate signaling.

### Exposure to ethanol has no effect on AMPA/NMDA ratio

Changes in AMPA/NMDA ratio has been considered a hallmark of learning and memory processes and drugs of abuse can ‘hijack’ these processes to cause maladaptive plasticity (Lüscher and Malenka, 2011). To answer how ethanol conditioning affected postsynaptic glutamatergic transmission in vBNST neurons. To test this, we measured AMPA-receptor mediated EPSCs by voltage clamping the cell at −70 mV and NMDA-receptor mediated currents by holding the same cell at +40 mV. This allows us to calculate AMPA/NMDA ratio for each cell (see methods). We found no significant differences in AMPA/NMDA ratio across all the different experimental groups (Fig. 5G-I; [t(35)=0.0132, p=0.9896] in CPP group; [t(37)=0.2002, p=0.8424] in CPA group; [t(18.06)=0.5317, p=0.6014; unpaired t-test with Welch’s correction] in Home cage group).

## Discussion

Cue-induced drug seeking behavior has been known to contribute to relapse from abstinence in substance use disorders (Anton, 1999; Bardo and Bevins, 2000). When the rewarding properties of drugs become associated with environmental cues, a previously neutral cue acquires salience through a form of Pavlovian conditioning and can trigger drug seeking. Drugs of abuse are complex pharmacological compounds that produce both positive rewarding and negative aversive properties. Prior work from Cunningham and colleagues has demonstrated that the motivational effects of alcohol varied as a function of time after injection (Cunningham *et al*, 1997; Cunningham and Henderson, 2000). Specifically, ethanol injection produces an initial short-duration aversive effect followed by a longer-lasting rewarding effect. The same group has also shown that pre-exposure to ethanol before place conditioning resulted in tolerance to its aversive effect without modifying the rewarding effect (Cunningham *et al*, 2002). Therefore, it is imperative to take into consideration the balance between reward and aversion when trying to model drug seeking behavior.

In the present study, we utilized a well characterized form of Pavlovian conditioning to assess the rewarding as well as aversive properties of ethanol (for review see Tzschentke, 1998) and coupled that with ex vivo patch clamp electrophysiology to determine the effects of ethanol conditioning on synaptic plasticity in the BNST. To our knowledge, this is the first study to look at electrophysiological correlates of both ethanol-induced CPP and CPA in the BNST. We focused on ventral BNST, as it is enriched with projection neurons and plays a key role in addiction-related behavior. We found that the vast majority of vBNST neurons had high input resistance (typically > 400 MΩ) and low whole-cell capacitance (typically < 30 pF) along with a smaller subpopulation of neurons with low input resistance (typically between 100-300 MΩ). Prior work from our lab and others have shown that high input resistance and low capacitance neurons tend to be associated with putative vBNST-VTA projecting neurons (Dumont and Williams, 2004; Kash et al, 2007). We found an increase in intrinsic excitability of vBNST neurons following ethanol conditioned preference without any changes in synaptic transmission. In contrast, our data from mice that showed ethanol conditioned aversion indicate that synaptic transmission onto vBNST neurons is significantly altered with a marked decrease in spontaneous glutamatergic transmission resulting in excitatory-inhibitory imbalance. These results provide evidence for distinct neuroadaptations in the BNST induced by contextual learning associated with both rewarding and aversive properties of ethanol. Together, our data add to the current literature supporting the role of BNST in reward seeking behavior and drug addiction.

### Ethanol-evoked plasticity in vBNST neurons following conditioned preference

We found an increase in neuronal excitability of vBNST neurons following expression of ethanol-induced CPP. The increase in intrinsic excitability of these neurons was associated with lowering of the threshold for action potential initiation resulting in greater spike frequency in response to current injection. This change in excitability was not observed in mice that received ethanol injections in their home-cages (unpaired) suggesting that presentation of ethanol associated-cues can drive hyperexcitability of vBNST neurons distinct from the pharmacological effects of ethanol alone. Furthermore, we observed a decrease in latency to fire action potentials, which can contribute to increased firing and may be induced by plasticity of various ion channels, particularly potassium channels which are known to be key regulators of neuronal excitability (Lüscher and Slesinger, 2010; Kourrich *et al*, 2015).

A family of potassium channels known as inwardly-rectifying potassium channels often reduce neuronal excitability by hyperpolarizing the neuron through the entry of potassium ions near resting membrane potentials (Matsuda *et al*, 1987). Inwardly rectifying potassium channels are activated by neurotransmitters and hormones as well as by other messengers such as kinases and G-proteins (Dascal, 1997). The G protein-activated potassium (GIRK) channels can be directly activated by ethanol (Aryal *et al*, 2009) and have been found to play a role in drug addiction (Herman *et al*, 2015; McCall *et al*, 2016; Munoz *et al*, 2016; Rifkin *et al*, 2017). Here we show that ethanol-induced CPP increases the excitability of vBNST neurons by potentially blocking inwardly-rectifying potassium channels. Tertiapin-Q (200 nM), a selective blocker of inward-rectifier potassium channels, increased the excitability of vBNST neurons in saline-treated mice but did not alter the firing properties of these neurons in ethanol-treated mice. Interestingly, the decrease in latency to fire AP as well as the lowered threshold for firing observed in the EtOH-CPP mice compared to Sal-CPP mice still persisted in the presence of TPQ. This would suggest the concurrent involvement of a different potassium channel (such as A-type or D -type) in regulating the neuronal excitability in vBNST. Our findings contribute to a growing body of literature implicating inward-rectifier potassium channels in alcohol abuse and the rewarding effects of alcohol. For example, global knock-out of a subunit of GIRK channel (GIRK3) resulted in enhanced ethanol-CPP (Tipps *et al*, 2016) but knocking out GIRK2 subunit reduced ethanol-induced CPP (Hill *et al*, 2003).

Overall, similar changes in intrinsic excitability following cocaine conditioning have been observed in other brain regions such as prefrontal cortex (Dong *et al*, 2005), laterodorsal tegmental nucleus (Kamii *et al*, 2015) and nucleus accumbens (Kourrich and Thomas, 2009). Chronic exposure to ethanol also results in increased excitability of vBNST (Marcinkiewcz *et al*, 2015; Pleil *et al*, 2015). In juxtacapsular BNST, high frequency stimulation results in long-term potentiation of intrinsic excitability which was impaired following protracted withdrawal from self-administration of alcohol (Francesconi *et al*, 2009). Therefore, increases in intrinsic excitability may provide a permissive function resulting in increased temporal fidelity of firing thereby predisposing the neural network to generate long-term neuronal changes.

In contrast to intrinsic plasticity, we did not observe any effects of ethanol conditioning on both presynaptic and postsynaptic transmission. There were no changes in either frequency or amplitude of excitatory or inhibitory synaptic transmission. Neither did we observe any changes in the AMPA/NMDA ratio. It is possible that the short-term increase in intrinsic excitability induces long-term homeostatic adaptations in synaptic strength that develops later than the timescale at which we recorded from the neurons. An alternate rationale for the lack of effect of ethanol-CPP on vBNST synaptic transmission is the cell-type and projection-target heterogeneity of BNST neuronal populations. In dorsal BNST, Kim *et al*, 2013 found that the oval nucleus and the anterodorsal BNST had divergent effects on anxiety state. In a different study, VTA-projecting glutamate and GABA neurons had opposite effects on anxiety and motivated behavior (Jennings *et al*, 2013). Also, prior work from our lab demonstrated the differential modulation of interneurons and projection neurons in regulating emotional behavior (Marcinkiewcz *et al*, 2016). There is also evidence for discrete genetically defined neuronal populations being modulated by alcohol (Silberman *et al*, 2013; Pleil *et al*, 2015b). Thus, future work will need to target discrete cell- and projection-specific neuronal populations to capture more nuanced changes in synaptic plasticity and also layer on how various neuromodulators that are involved in alcohol-induced plasticity (for example CRF, NPY and serotonin, dopamine) can affect this form of plasticity.

Altogether, these findings support the broad literature implicating the role of BNST in reward seeking and drug-associated behavior (Lovinger and Kash, 2015; Vranjkovic *et al*, 2017). Specifically, context-associated ethanol-seeking results in increased c-Fos immunoreactivity (Hill *et al*, 2007), and, chemogenetic inactivation of BNST in either a non-specific manner (Pina *et al*, 2015) or selective inhibition of BNST-VTA projection neurons (Pina and Cunningham, 2017) can attenuate the expression of ethanol CPP. Our data offer additional insight by providing a neural framework underlying ethanol-seeking behavior and the involvement of BNST in drug-associated memories.

### Neuroadaptations in vBNST neurons following ethanol-conditioned aversion

BNST is an integral hub not just for reward-related behavior but for regulating aversive learning and anxiety-like states (for review, see Lebow and Chen, 2016; Vranjkovic *et al*, 2017). It is involved in a number of behaviors relevant to aversion and anxiety-like states, such as, the acquisition and expression of Pavlovian fear conditioning (Lebow and Chen, 2016), stress-induced reinstatement of drug seeking (Mantsch *et al*, 2015), and, negative affect in pain (Minami, 2009). Using a conditioning paradigm that has been shown before to cause ethanol-induced conditioned aversion (Cunningham *et al*, 1997, 2006), we looked at synaptic alterations in vBNST neurons. Conditioned aversion caused significant changes in presynaptic plasticity that were distinct from the changes following conditioned preference. Unlike ethanol-CPP, ethanol-CPA didn’t induce changes in neuronal excitability when compared to respective saline controls. Our method of examining synaptic transmission allows us to measure both excitatory and inhibitory transmission from each individual neuron to directly assess the balance of excitatory and inhibitory synaptic drive onto individual neurons. Using this approach, we observed a significant reduction in the frequency of spontaneous excitatory synaptic transmission following ethanol-conditioned aversion without altering the frequency of spontaneous GABAergic transmission. This resulted in an overall decrease in excitatory synaptic drive onto vBNST neurons. Interestingly, in the presence of TTX, which blocks action potential dependent synaptic transmission, this decrease in excitatory synaptic drive was lost. These results suggest an activity-dependent modulation of glutamate signaling. Our hypothesis was further bolstered by changes in short-term plasticity of evoked glutamatergic responses that resulted in an increase in synaptic facilitation. The remodeling of glutamatergic synapses have been postulated to play a role in classical learning and memory (Malenka and Bear, 2004) and although we show compelling evidence for short-term plasticity at glutamatergic signaling following ethanol-conditioned aversion, the source(s) of glutamate as well as the underlying neural mechanism that is being modified in response to conditioning is unclear.

Within BNST, modulation of glutamatergic transmission in response to stress and drug seeking behavior is complex and involves both postsynaptic and presynaptic metabotropic receptors and is modulated by various neuropeptides and monoamines (for extensive review, see McElligott and Winder, 2009; Harris and Winder, 2018). One likely candidate for modulation of both short-term and long-term depression at glutamatergic terminals within the BNST is the endocannabinoid signaling which acts retrogradely to inhibit release probability at presynaptic terminals (Grueter *et al*, 2006; Puente *et al*, 2011). Recently, we showed that Gq-mediated signaling in the GABA transporter (VGAT)-expressing neurons promotes anxiety-like behavior that was accompanied by long-term plasticity at CB1R-dependent glutamate signaling (Mazzone *et al*, 2016b). Another equally potential candidate is the noradrenergic input to the vBNST, which is also important for mediating negative affect and anxiety-like behavior. Specifically, disruption of noradrenergic signaling in the BNST attenuates opiate-withdrawal-induced CPA (Delfs *et al*, 2000) and also formalin-induced CPA (Deyama *et al*, 2008) and induces plasticity at glutamatergic terminals (Egli *et al*, 2004; McElligott *et al*, 2010). Further studies are warranted to parse out the mechanism of modulation of various glutamatergic inputs onto vBNST following ethanol-conditioned aversion. Potentially, the decreased excitatory drive that we observed may act as a filter that allows only strong and salient synaptic inputs to activate the neuron, thereby enhancing the signal-to-noise ratio.

### Functional implications

While ethanol-induced conditioned place preference results in increased intrinsic excitability of vBNST neurons; ethanol-induced conditioned place aversion selectively modulates glutamate signaling to reduce excitatory synaptic drive onto vBNST neurons. These BNST-specific neuroadaptations likely alter the activity dynamics of distinct BNST outputs. One such output is BNST-VTA projection, which plays a complex role in modulating both aversive and rewarding behaviors (see: Jennings *et al*, 2013; Glangetas *et al*, 2015; Marcinkiewcz *et al*, 2016; Pina and Cunningham, 2017). BNST-VTA projections are thought to preferentially innervate non-dopaminergic VTA neurons (Jennings *et al*, 2013), and can indirectly increase the activity of dopamine neurons by disinhibiting GABAergic neurons in the VTA. Thus, it is likely that ethanol conditioning alters BNST to VTA circuit to regulate both rewarding and aversive learning. Collectively, in this study we have presented compelling evidence for neural plasticity in the ventral part of BNST following both ethanol-conditioned preference and aversion. This provides a conceptual framework for future experiments to identify specific mechanisms that contribute to both these forms of ethanol-induced associative learning and their role in overall regulation of ethanol seeking behavior that contributes to relapse.

